# The CCR2 inflammatory pathway is a target for improving severe disease and pulmonary inflammation in experimental COVID-19

**DOI:** 10.1101/2025.03.28.645941

**Authors:** Yvette Kazungu, Shrilakshmi Hegde, Parul Sharma, Amy Marriott, Andrew Steven, Jesus Reine, Jessica Dagley, Matthias Mack, James Stewart, Joseph D Turner

**Affiliations:** Centre for Drugs & Diagnostics, Centre for Neglected Tropical Diseases, Department of Tropical Disease Biology, Liverpool School of Tropical Medicine, Liverpool, United Kingdom; Institute of Infection, Veterinary & Ecological Sciences, Faculty of Health and Life Sciences, University of Liverpool, Liverpool, United Kingdom; Department of Internal Medicine II – Nephrology, University Hospital Regensburg, Regensburg, Germany

## Abstract

SARS-CoV2 can induce an acute respiratory distress syndrome (ARDS), provoked by a dysregulated hyper-inflammatory pulmonary immune response. Here we used the keratinocyte-18 humanized angiotensin converting enzyme-2 (K18-hACE2) mouse model of SARS-CoV2, where expression of the CoV2 spike protein receptor, hACE-2, is restricted to epithelia, to characterize inflammatory pulmonary immune responses post-intranasal infections with the delta isolate SARS-CoV2(Δ B.1.617.2). Immune-profiling by focused transcript analysis, inflammatory protein array, and multi-color flow cytometry, confirmed that clinically relevant markers of COVID-19 (IL-6, GM-CSF, neutrophils, inflammatory monocytes) were significantly elevated in lungs of mice at day 5 post-infection and that remdesivir anti-viral active metabolite (GS441524) treatment significantly modified SARS-CoV2Δ viral loads and pulmonary-specific inflammation. Chemokine ligands of CCR2 (CCL2/7/8) were amongst the top 5% upregulated pulmonary transcripts in a focused human infection response array to SARS-CoV2Δ. CCL2 was confirmed as elevated in protein assays in SARS-CoV2Δ infected lungs. To address functional relevance of the CCR2 pathway of inflammatory cell recruitment to the lungs mediating disease, mice were administered with anti-CCR2 antibody daily at point of infection for up to six days. Anti-CCR2 treated mice showed significant improved welfare scores, were protected from weight loss, modified myeloid pneumonitis, and displayed significantly blunted cytokine and chemokine response in the lungs, despite not affecting pulmonary viral loads. Our data supports therapeutic benefit of modifying CCR2-dependent cell recruitment in the treatment of viral-induced ARDS.

## Introduction

Acute Respiratory Distress Syndrome (ARDS), a severe and life-threatening manifestation of acute lung injury and respiratory failure, characterized by pneumonitis and hypoxemia, is triggered by a variety of infectious and non-infectious causes, including sepsis, cigarette smoking, Immunotherapeutics, vaping and viral infections^1^. Periodic viral infections of pandemic potential such as H1N1 swine flu, MERS and SARS, are recent major emergent drivers of ARDS leading to hospitalization, with potential for subsequent multi-organ failure and mortality in vulnerable patient populations. Over 7 million SARS-CoV2 deaths have been reported to the World Health Organization as of March 2025^2^. Given the high social and economic burdens of lung virus disease, there is an urgent need to increase our understanding of the aetiology of ARDS toward identifying novel therapeutic targets which may be modified with effective treatments to limit severe morbidity and mortality.

Respiratory viral infections mediating epithelial tissue damage can subsequently trigger a complex cascade of deleterious hyper-inflammatory immune responses within the lungs which, if not appropriately regulated, lead to exacerbated lung injury^3^. SARS-CoV2 clinical severity grading including ARDS, oxygen administration, hospitalization, mechanical ventilation, and mortality, is significantly and progressively associated with elevated systemic cytokines / chemokines (so-called cytokine storm), particularly CXC ligand (CXCL)-10, interleukin (IL)-6, and Granulocyte Macrophage Colony Stimulating Factor (GM-CSF)^4^. This inflammatory milieu is thought to contribute to the expansion of blood leukocytes and their recruitment and activation within pulmonary tissues. For example, late-stage severe COVID-19 is associated with neutrophilia, and in autopsy patients, an influx of monocytes/macrophages into the lung parenchyma and a myeloid pulmonary artery vasculitis have been determined^5,6^. Dysregulated myeloid infiltrates into pulmonary tissues are proposed to induce a positive feedback loop via cytokine and chemokine signaling for cumulative intra-pulmonary cellular trafficking. Myeloid infiltrates contribute to a progressive ‘immuno-thrombotic’ state, whereby release of neutrophil DNA extracellular traps (NETs) and clotting factor release by activated monocytes, such as Tissue Factor, contribute to micro-thrombi formation in the pulmonary vasculature^3^.

Targeting facets of this pro-inflammatory cascade with specific anti-inflammatory treatments is thus a promising therapeutic approach to prevent ARDS-related mortality. Indeed, the large-scale multi-centre phase III trial platform: Randomised Evaluation of COVid-19 thERapY (RECOVERY), has established that the steroidal anti-inflammatory, dexamethasone, the Janus kinase (JAK) inhibitor barcitinib, and the interleukin-6 receptor antagonist, tocilizumab are efficacious live saving treatments for hospitalized cases of SARS-CoV2 related ARDS^7–9^. The keratin18 (K18)-human angiotensin converting enzyme (ACE)2 mouse model was originally developed as a susceptible laboratory disease model of the SARS-1 Coronavirus^10^. Expression of the human ACE2 receptor in the lower airway epithelium enables the transition from mild to severe disease in mice following exposure to SARS-CoV2, and a more comprehensive disease profile recapitulating severe ARDS human cases. This has enabled the use of K18-hACE2 mice as a preclinical screen to interrogate novel therapeutic interventions against SARS-CoV2, including antiviral small molecules and antibody-based therapeutics^11–15^.

Here, we utilized experimental infections of K18-hACE2 mice with the highly pathogenic SARS-CoV2 Delta variant of concern (VoC; B.1.617.2)^16^ and report the lung-specific immune profiles during infection using an array of pro-inflammatory multiplex immunoassay and host-response gene expression panels. We determine several C-C Motif Chemokine Receptor 2 (CCR2) chemokines are amongst the most abundant pulmonary-specific markers of SARS-CoV2 delta infection, are modified effectively by remdesivir metabolite anti-viral therapy, and, by use of a CCR2 neutralizing antibody, show the benefit of specifically targeting CCR2 host responses in ameliorating pulmonary cytokine storm and acute clinical disease.

## Materials and Methods

### Ethics statement

All infection study protocols were approved by Animal Welfare and Ethical Review Bodies from University of Liverpool and Liverpool School of Tropical Medicine. All animal work was performed in accordance with the UK Home Office requirements (Home Office licence: PP4715265). All risk assessments and protocols were pre-approved by the relevant authorities and infection studies were carried out in containment level 3 by trained personnel.

### In vivo mice infection studies

Male and female heterozygous K18-hACE2 c57BL/6J mice (strain: B6.Cg-Tg(K18-ACE2)2Prlmn/J) aged 6-8 weeks old were purchased from Charles River Laboratories and maintained under SPF barrier conditions in individually ventilated cages. Animals were randomly assigned to different experimental groups per cage and fed a standard chow diet. All infections reported herein used the B.1.617.2 (Delta variant) hCoV-19/England/SHEF-10E8F3B/2021 (GISAID accession number EPI_ISL_1731019). Mice were inoculated intranasally with 1000-2000PFU in 50µl of normal saline under light anesthesia. Mock infected animals were exposed to 50ul of saline intranasally.

The active metabolite of the antiviral Remdesivir (GS441524; Tocris bioscience, Bristol, UK) was administered intraperitoneally (ip) once daily at 50mg/kg, as described previously^17^. The rat anti-mouse CCR2 monoclonal antibody (MC-21)^18–20^ was administered ip at 20µg in 100µl normal saline. Mock controls received 100 µl normal saline. Because of strict operational conditions at high containment labs, treatment groups were not blinded during experiments. All animals were weighed daily and monitored for welfare signs two times daily. Animals were evaluated for changes in appearance, natural behavior, provoked behavior, food/water intake and respiration. They were graded 0 to 3 (0-none, 1-mild, 2-moderate, 3-severe). Following 4 – 6 dpi, animals were humanely euthanized using approved method (rising concentration CO_2_ followed by confirmation of cardiac arrest) and blood, nasopharyngeal swabs and lung tissues were collected for downstream processing.

### Flow Cytometry

Lung tissues were digested with 500μl HBSS + 0.1 mg/mL DNase (Roche) +0.25 mg/ml Liberase TL (Merck), 37°C, 45 minutes. A further 500 μl of 0.5% 2mM EDTA, 2% FBS in PBS was added and samples incubated for 5 minutes, 37°C. Digests were sieved (40 μm sieve) before centrifugation for 5 minutes at 4°C and 400xg. Cell pellets were resuspended in 1 ml RBC lysis buffer (Invitrogen) and incubated at room temperature for 3 mins. RBC lysis was neutralized by 10x volumes of PBS and pelleting cells by centrifugation (400xg for 5 minutes). Cells were resuspended in 0.1ml FACS buffer (BD Biosciences). Single-cell suspensions were stained with viability dye followed by specific monoclonal antibodies against specific markers for 25 min at 4°C at dilutions listed (Table S1). Cells were washed and fixed with 1X BD CellFix (BD Biosciences). Single stained compensation controls were prepared using UltraComp ebeads (Thermo Fisher Scientific). Samples were acquired on a spectral cytometer (Cytek Aurora 4L 16UV-16V-14B-8R) with a gating strategy (SFig.1) prior reported ^21^.

### RNA extraction and qRT-PCR

Lung samples were homogenized in 500µl of TRIzol (Thermo Fisher Scientific) using TissueLyser LT (Qiagen). Total cellular RNA was extracted using the TRIzol manufacturer’s protocol. RNA samples were resuspended in nuclease-free water and quantified using an Implen Nanophotometer. A one-step quantitative real time (RT)-qPCR was performed to quantify viral RNA levels according to an optimized protocol^22^ in a 20μl reaction mix using the GoTaq® Probe 1-Step RT-qPCR System (Promega) utilising 500ng of RNA template. Primer/probe mixes used for nCOV_N1, E sgRNA, and murine 18s were purchased from Integrated DNA Technologies (listed in Table S2). Standard curves were generated with 10-fold serial dilutions of linearized standard plasmids from 10^8^ to 10^1^ copies per reaction. All samples and standards were run in duplicates using the following RT-qPCR cycles; 45°C for 15 minutes, 95°C for 2 minutes, 95°C for 3 seconds, and 55°C for 30 seconds for 44 cycles.

### Plaque assay

Lung tissue samples were homogenized in DMEM using TissueLyser LT (Qiagen) and supernatants were collected. Vero-E6 cells (ATCC) were cultured in complete medium (Dulbecco’s modified Eagle’s medium (DMEM) supplemented with 5% complement deactivated FBS) at 37°C / 5% CO_2_. Vero E6 cells were seeded in 24-well plates (100000 cells/well) and left to grow overnight. Cell monolayers were infected in duplicate with 10-fold serial dilutions of SARS-CoV-2 supernatants in serum free Minimum Essential Medium (MEM) for 1 hour incubation at 37°C, 5% CO_2_. A 1ml overlay containing 2% Agarose in complete medium (1:4 ratio) was added and incubated further for 3 days. Cells were fixed *in situ* with an addition of 4% PFA solution (Thermo Scientific) 30 minutes at room temperature. Monolayers were stained with 1% crystal violet solution (Sigma Aldrich) and incubated for 20 minutes at room temperature. Plates were washed with water until plaques were visible for enumeration.

### Protein and immuno-assays

Lung samples were homogenized in T-PER lysis buffer (Sigma Aldrich) with protease inhibitor cocktail at 1:100 (Sigma Aldrich). Lysates were clarified at 10000xg for 15 minutes at 4°C and supernatants were then quantified using the Pierce BCA protein assay kit (Thermo Fisher Scientific) according to the manufacturer’s instructions. 1000ng of lung tissue homogenates was used for the analysis of cytokines and chemokines by MCYTMAG-70K-PX32 and MTH17MAG-47K-12C panels (Millipore) according to the manufacturer’s instructions. The assays were then run on the Luminex 200 platform utilizing a 5PL curve fitting model to calculate analyte concentrations.

### NanoString gene expression analysis

A master mix containing hybridization buffer and the reporter code set was aliquoted (8μl) and 5μl containing 100ng of target RNA and 2μl of the capture probe set were added. This reaction was incubated at 65°C, overnight to complete hybridization. A digital analyser scanned and counted the molecular barcode per sample at 555 fields of view (FOV). Data was analyzed by ROSALIND® (https://rosalind.bio/), with a HyperScale architecture developed by ROSALIND, Inc. (San Diego, CA). Gene expression was quantified by fluorescent signals, determined by the level of probe-RNA hybridizations^23^. Normalization, fold changes and p-values were calculated using criteria following the nCounter Advanced analysis protocol provided by <u>NanoString</u>.

### Statistical analysis

Statistical analyses were carried out using GraphPad Prism 9 (GraphPad, Inc., La Jolla, CA). All continuous data were tested for normal distribution using the Shapiro wilk test. Where data were normally distributed, a 2-tailed independent Student’s t-test (2 groups) or 1-way ANOVA test (>2 groups) was used with Tukey or Sidak’s post-hoc test for significant differences. To compare two independent variables between the 2 or more groups with normal distribution, 2-way-ANOVA was used with Dunnett’s post-hoc test. Where data were found to be not normally distributed, a two-tailed Mann-Whitney U test or Kruskal-Wallis with Dunn’s post hoc multiple-comparison test (>2 groups) was utilized to test significant differences between groups.

Survival analysis was performed using the Kaplan-Meier method. Significant p-values are denoted by asterisks (p≤0.05 = *, p≤0.01=**, p≤0.001 = *** and p≤0.0001 = ****).

## Results

### SARS-COV2Δ infected K18-hACE2 mice recapitulate clinical severe disease with upregulation of multiple pulmonary-specifc inflammatory markers

We evaluated infectivity, progression to severe disease and subsequent associated pulmonary cellular inflammatory responses to SARS-CoV2Δ post-intranasal experimental infection in K18-hACE2 mice (Fig.1A). Initial studies with both male and female mice revealed that male mice were more susceptible to infection and developed severe pulmunary disease than their female counterparts when compared in weight changes, welfare scores and viral loads (SFig. 2). Subsequently, we limited experiments on male mice to discover abundent inflammatory markers of severe clinical disease.

**Figure 1:**
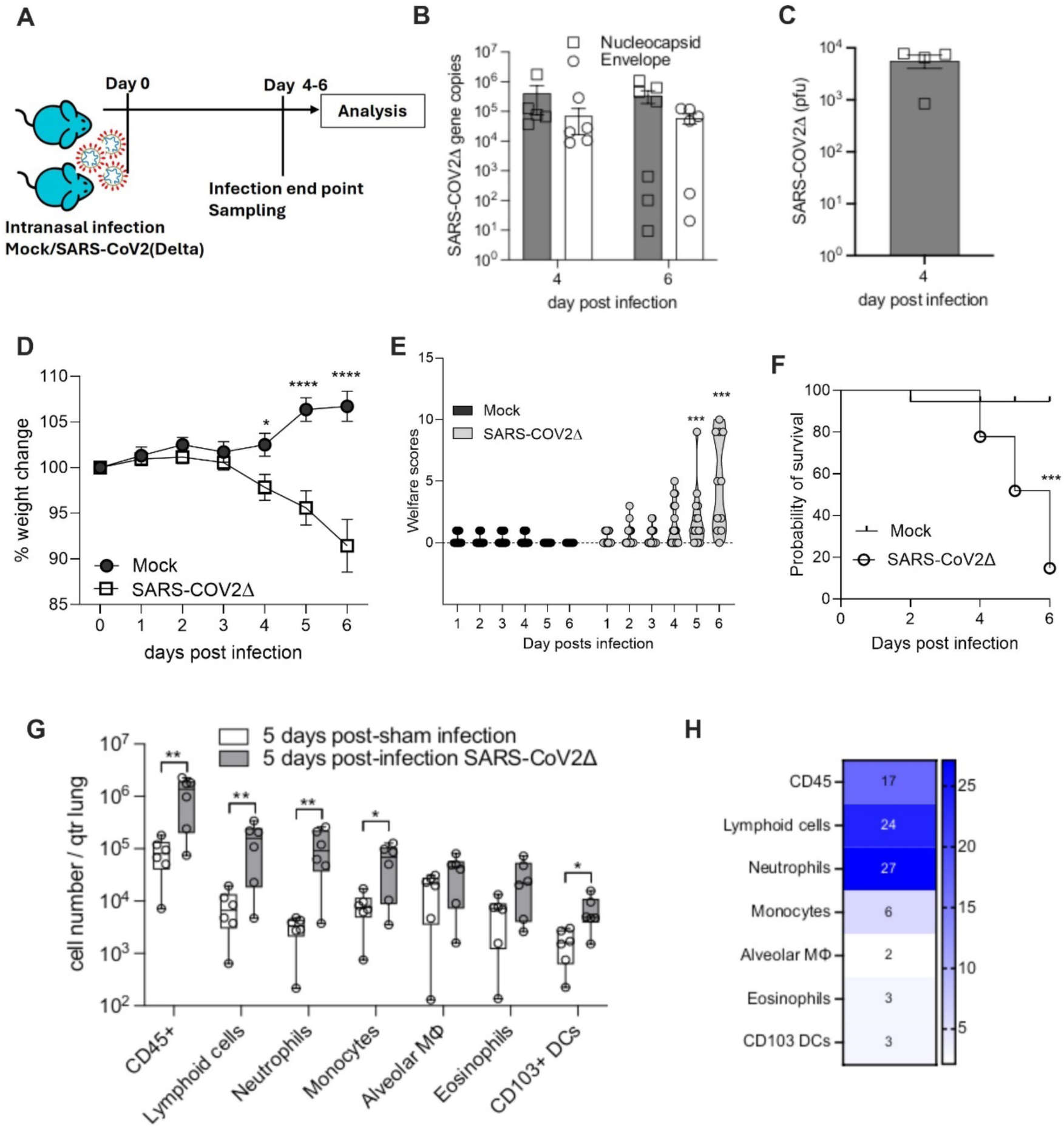
SARS-CoV2Δ-mediated acute severe cellular inflammatory lung disease is induced in K18-hACE2 mice. **A**. Experimental design showing intranasal infection in with 1000-2000 PFU/ml of SARS-CoV2 (Δ-Delta variant) or normal saline (Mock) and observed for up to 6 days. **B**. One step qPCR quantification of N1 and sgE RNA copies in lung tissue at 4 and 6 dpi and **C**. viral yields from lung tissues by plaque assay at 4 dpi. Data for individual mice are overlayed onto histograms representing mean±SEM per group. **D**. Weight changes and **E**. welfare scores in SARS-CoV2 infected mice compared with mock controls 1-6 dpi. Data represented as mean±SEM (p≤0.05 = *; p≤0.001 = ***; p≤0.0001 = ****; n=8-10 / group). **F**. Cumulative survival probability of 18 mice over 6 days of infection across 3 experiments (log-rank analysis, p≤0.001 = ***; n=8-10 / group). **G**. Box plots showing absolute cell counts of different immune cell sub-types in mock and SARS-CoV2 infected lungs. Each dot represents an individual animal. (multiple unpaired t-test, p≤0.05 = *; p≤0.01 = **). H. Increase in immune cells sub-types in SARS-CoV2 infected lung represented as median fold change compared to mock controls (n=6 / group).

Reproducible infections at 4 and 6 days post infection (dpi) were determined in lung tissues by qPCR quantification of both nucleocapsid (N1) and subgenomic E (sgE) protein RNA copies as well as viral yields quantification by plaque assay (Fig.1B, 1C). We observed significant weight loss in infected mice compared to mock control group from 4dpi (Fig.1D; p<0.05, mixed-effect 2-way ANOVA, Šídák’s multiple comparisons post-hoc test). Also, welfare scores were significantly higher in infected mice from 5dpi onwards, suggesting decrased overall health (Fig.1E; p<0.05, Mann-Whitney multiple comparison test). Cumulative data of 18 mice assessed across 3 experiments showed significant decrease in survival by 6dpi, defined by attaining a theshold of clinical severity and/or weight loss of ≥10% that mandated a humane endpoint for the viral infection (Fig.1F; 28% survival probability, 5/18 mice assessed, p=0.0005, log-rank analysis for survival).

To characterise inflammatory cells in SARS-CoV2Δ infected lungs, we immunophenotyped and compared different immune cell lineages between mock and SARS-CoV2Δ infected groups using a previously published flow cytometry panel, focusing on myeloid pulmonary infiltrates following lung injury^21^. Following 5dpi, we found a significant increase in the recruitment of immune cell populations associated with hyper-inflammation and severe disease in lung tissues (Fig 1G-H,). Specifically, total CD45^+^ leukocytes counts were significantly increased by average (median) 17-fold in infected lungs, a significant increase in MHC-II^+^ CD11b^low^ lymphoid cells (median fold change = 24), CD11b^+^ Ly6G^+^ neutrophils (median fold change = 27), CD11b^+^ Ly6C^+/−^ monocytes (median fold change = 6), and CD11b^−^ CD103^+^ CD24^+^ CD11c^+^ dendritic cells (median fold change=3) in infected group compared to mock infection controls (Fig 1G-H; p<0.05, multiple unpaired t-test). The lung pneumonitis profile identified here closely aligns with available data from clinical lung samples following SARS-CoV2Δ infection, thus further validating the K18-hACE2 mice infection model as an appropriate disease model for testing new therapeutic approches against the pathogen.

### Remdesivir GS441524 treatment of infected K18-hACE2 mice limits SARS-CoV2**Δ** replication and modifies inflammatory markers associated with severe lung disease

Having initially established the profile of SARS-CoV2Δ K18-hACE2 cellular lung inflammation, we subsequently utilised the model to evaluate effect of anti-viral treatment on lung pathology utilising the active metabolite of Remdesivir, GS441524, at 50mg/kg (ip)^17^ (Fig. 2A). Remdesivir is a broad-spectrum antiviral that has been used against multiple viruses^24^. Although efficacy of remdesivir and GS441524 in clearing SARS-CoV2 viral load has been previously documented^25–30^, its effect on lung pathology and inflammation reduction has not been studied. Our data shows that GS441524 treatment significantly reduced both N1 and sgE viral RNA transcripts in nasopharyngeal cavity at 4 days post infection (p≤0.05 = *; two-tailed unpaired t-test, Fig. 2B-C) and in viral yields from lung tissue supernatants as quantified by plaque assay (p≤0.05 = *; two-tailed unpaired t-test, Fig. 2D).

**Figure 2:**
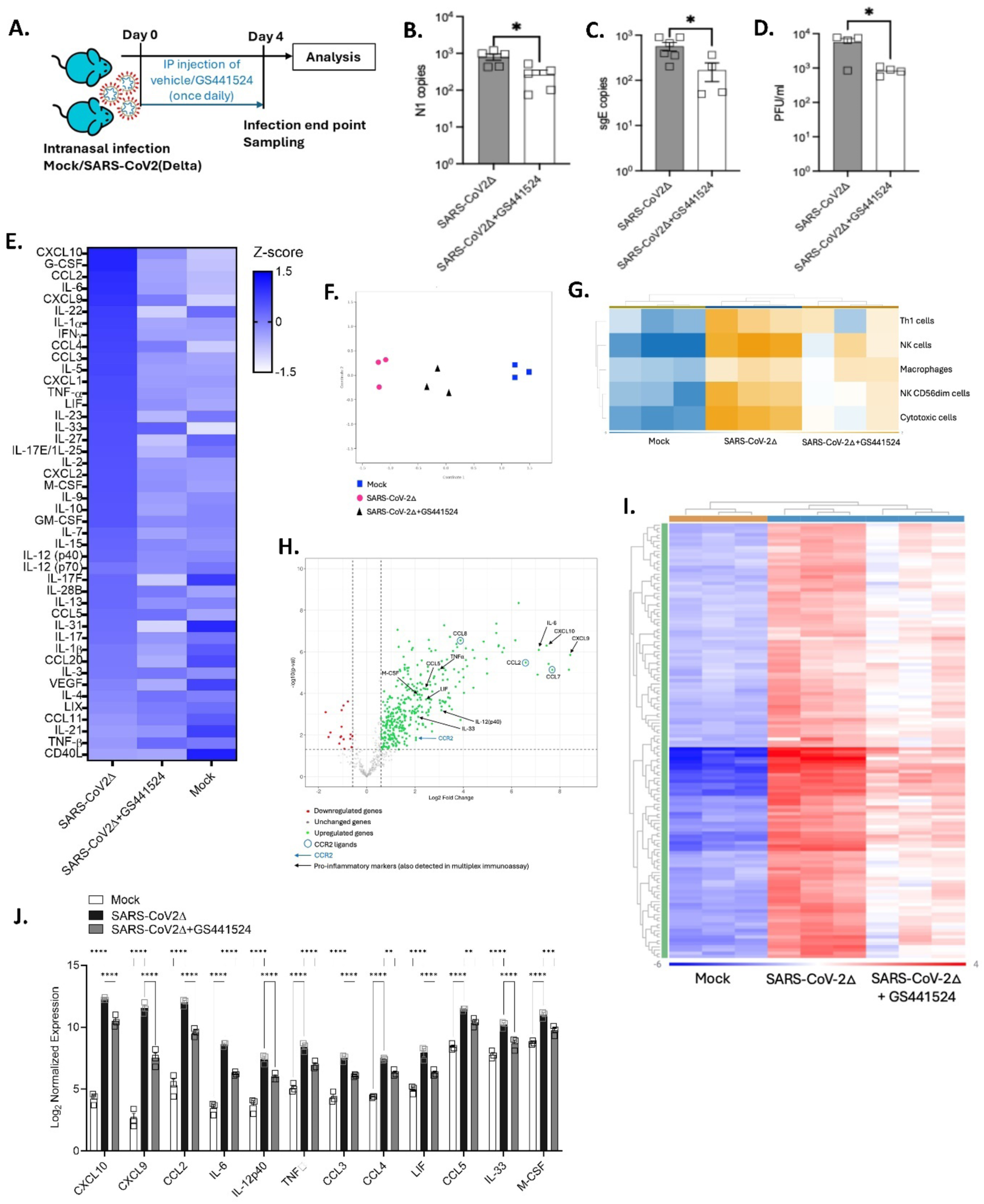
Remdesivir active metabolite GS441524 anti-viral treatment of SARS-CoV2Δ infected K18-hACE2 mice reduces viral load and decreases lung inflammation. **A.** Mice (n=6 per group) were infected with 2000 PFU of SARS-CoV2Δ or normal saline (Mock), a group of infected mice also received treatment with GS441524 for the duration of the experiment. **B.** N1 **C.** sgE viral RNA copies from nasopharyngeal swabs quantified using one-step qPCR and **D.** viral yield quantification from lung tissues by plaque assay. Data for individual mice are overlayed onto histograms representing mean±SEM per group (p≤0.05 = *; two tailed, unpaired t-test). **E**. Heatmap showing z-scores of cytokines and chemokines in lung lysate quantified by a multiplex immunoassay. (n=4-5/group) F. Nanostring analysis of a 785 gene mouse host response panel, multidimensional scaling showing distinct clusters among the experimental conditions **G**. Cell type profiling showing cell abundance scores for all samples **H**. Volcano plot showing upregulated and downregulated genes at fold changes −1.5 to 1.5, p-Adj=0.05 comparing infected and mock samples. **I.** Hierarchical clustering heatmap depicting the patterns of gene transcription between mock, SARS-COV2 infected and GS441524 treatment groups. **J**. List of significantly altered inflammatory markers following SARS-CoV2 infection that were identified in both transcriptomic analysis as well as in multiplex immunoassay. Data represented as mean±SEM. (p≤0.01 = **; p≤0.001 = ***; p≤0.0001 = ****; 2-way ANOVA with Dunnett’s post-hoc test).

To measure the effect of GS441524 on pro-inflammatory markers associated with SARS-CoV2Δ pulmonary ‘cytokine storm’, a multiplex immunoassay using a 44 cytokine/chemokine panel was performed. SARS-CoV2Δ infection invoked an aggressive pro-inflammatory response in lungs, with significant increase in multiple cytokines and chemokines such as CXCL10, G-CSF, CCL2, IL-6, CXCL9, CCL4, LIF, IL-33, CD-40L (Fig. 2E and SFig. 3, p<0.05, One way ANOVA, Tukey’s post-hoc test vs mock). We also observed relative increase in the levels of other cytokines and chemokines such as IL-1α, TNF-α, IFN-γ, IL-5, IL-10 and VEGF (p>0.05, Fig. 2E and SFig. 3). These results reflect elevated plasma cytokine levels reported in SARS-COV2 infected patients who presented with pneumonia which showed increase in multiple analytes including IL-1β, IL-7, IL-8, IL-9, IL-10, FGF, G-CSF, GM-CSF, IFN-γ, CXCL10, MCP-1, MIP-1α, MIP-1β, PDGF, TNF-α, and VEGF ^5,31–32^. Similar results have been reported in experimental mouse infection models including K18-hACE2 ^11,33^.

Further, we found that effective GS441524 treatment following the SARS-CoV2Δ infection concomitantly reversed the inflammatory phenotype by significantly decreasing IL-1α, IL-6, IL-22, IL-31, CXCL10, CCL2 and LIF levels compared to infected control group (p≤0.05, one-way ANOVA and Tukey’s post-hoc test, Fig. 2E and SFig 3). In addition, several other cytokines and chemokines including G-CSF, CXCL9, IFN-γ, CCL3, CCL4, IL-5, CXCL2, TNF-α, IL-25, IL-33, IL-2, CXCL1, M-CSF, IL-9, IL-10 and GM-CSF levels were relatively decreased following the GS441524 treatment (p>0.05, Fig. 2E, SFig. 3)

### Host response gene expression following SARS-CoV2**Δ** infection in K18-hACE2 mice and impact of remdesivir metabolite GS441524 treatment

To corroborate and extend immune inflammatory immune profiling, we undertook a comprehensive transcriptomic analysis in SARS-CoV2Δ infected lungs to characterize the in-depth inflammatory response following infection and simultaneously evaluated the impact of GS441524 treatment in K18-hACE2 mice. We utilized the ‘mouse host response gene expression panel’ from NanoString Technologies (Seattle, WA.), that quantifies approximately 785 genes and identifies significant pathways and host response dynamics using multiple available transcriptome data sets and pathway annotations. Gene set analysis showed multiple pathways were altered significantly following the infection compared to the mock control mice including those of pathogen signaling and inflammatory responses (SFig.4). Multidimensional scaling showed distinct clusters among the three experimental groups (Fig.2F) signifying unique individual group profiles. Cell type profiling analysis also showed relative increase in different inflammatory cells profiles in the infected group compared to mock/uninfected group. GS441524 treatment decreased the abundance of selected inflammatory cell types as compared to the vehicle treated, SARS-CoV2Δ infected group (Fig. 2G). Adjustment of p-values to account for multiple testing was performed using the Benjamini-Hochberg false discovery rate (FDR) method. Fold change cut-offs of ≥1.5 and ≤-1.5 and p-Adj=0.05 were used. A total of 337 genes were upregulated and 16 were downregulated while the rest remained unchanged in the infected lung compared to mock control. As we observed in protein immunoassay, several clinically relevant pro-inflammatory markers were upregulated (labelled and named in black, Fig. 2H). The chemokines and cytokines that were observed in both the multiplex immunoassay and gene expression analysis include CXCL10, CCL2, IL-6, CXCL9/MIG, IL-33 and LIF, many of these prior documented as characteristic of the cytokine storm observed following SARS-CoV2Δ infection ^5,31–32^. Additionally, heatmap analysis of gene expression among the three groups revealed unique transcriptome signatures. Both the uninfected and GS441524 treatment group displayed an overall decreased gene expression signature compared to the infected group (Fig. 2I). Normalized gene expression showed significant differences between the infected mice and those that were infected and treated with GS441524. Compared to uninfected mice lungs, there were only 189 genes that are significantly upregulated in the GS441524 treated mice lung following the infection, whereas 337 genes were upregulated in vehicle treated SARS-CoV2 infected mice lung. Specifically, we found 12 markers that were significantly increased in both multiplex immunoassay and transcriptomic profiling following SARS-CoV2Δ infection compared to mock infected lungs. Following the GS441524 treatment in infected mice, we found that all these 12 analytes were significantly decreased compared to vehicle treated infected group. (p<0.05, 2-way ANOVA with Dunnett’s post-hoc test, Fig. 2J). Altogether these results demonstrate GS441524 not only evokes an antiviral effect on SARS-CoV2Δ infection in the K18-hACE2 mouse model, but that effect also translates in the modification of the pro-inflammatory cytokine response that characterizes severe inflammatory lung disease.

### SARS-CoV-2**Δ** infection is characterized by an interferon stimulated gene (ISG) signature in K18-hACE2 mice which is preserved following remdesivir metabolite GS441524 treatment

Interferon signaling is critical in anti-viral innate immune responses and is a significant pathway upregulated during SARS-CoV-2 infection with an extensive Interferon Stimulated Genes (ISG) signature. ISGs including ISG15, MX1, IFIH1, IFIT2, IFIT3, Rsad2/Viperin, IRF7 have been recognized temporally in K18-hACE2 mice and human patients^11, 34–36^. In addition to interferon signaling, “RIG-I/MDA5 mediated induction of IFN α/β pathways” and “Immune system” were also identified as significantly altered pathways adding to the ISG signature that characterizes SARS-CoV2 infection (SFig 4). In this study, in addition to the above-mentioned genes, we have identified several other ISGs such as OAS3, IFIT1, IFI44, OASL1, OAS1A that are upregulated in the infected group (SFig 4). Interestingly, whilst GS441524 treatment reduced both the viral load and expression of pro-inflammatory cytokines and chemokines, ISG expression levels remained intact (SFig 4E).

### Targeting CCR2 ablates pulmonary hyper-inflammation and improves welfare in K18-hACE2 mice infected with SARS-CoV2**Δ**

Gene expression analysis of SARS-CoV2 infected lung identified CCR2 along with CCR2 ligands: CCL2, CCL7 and CCL8 as being some of the most overexpressed transcripts in SARS-CoV2Δ infected lungs (Fig. 2H, 3A). The CCL2-CCR2 pathway is postulated to be involved in the infiltration of CCR2-expressing ‘inflammatory’ monocytes into the lungs of patients following SARS-CoV2 infection, as CCL2 has been reported in the bronchoalveolar lavage fluid obtained from patients with severe COVID-19^37^. Evidence of an abundant pulmonary monocyte recruitment has been theorized to play a causal role in severe COVID-19 pathogenesis^38–39^.

**Figure 3:**
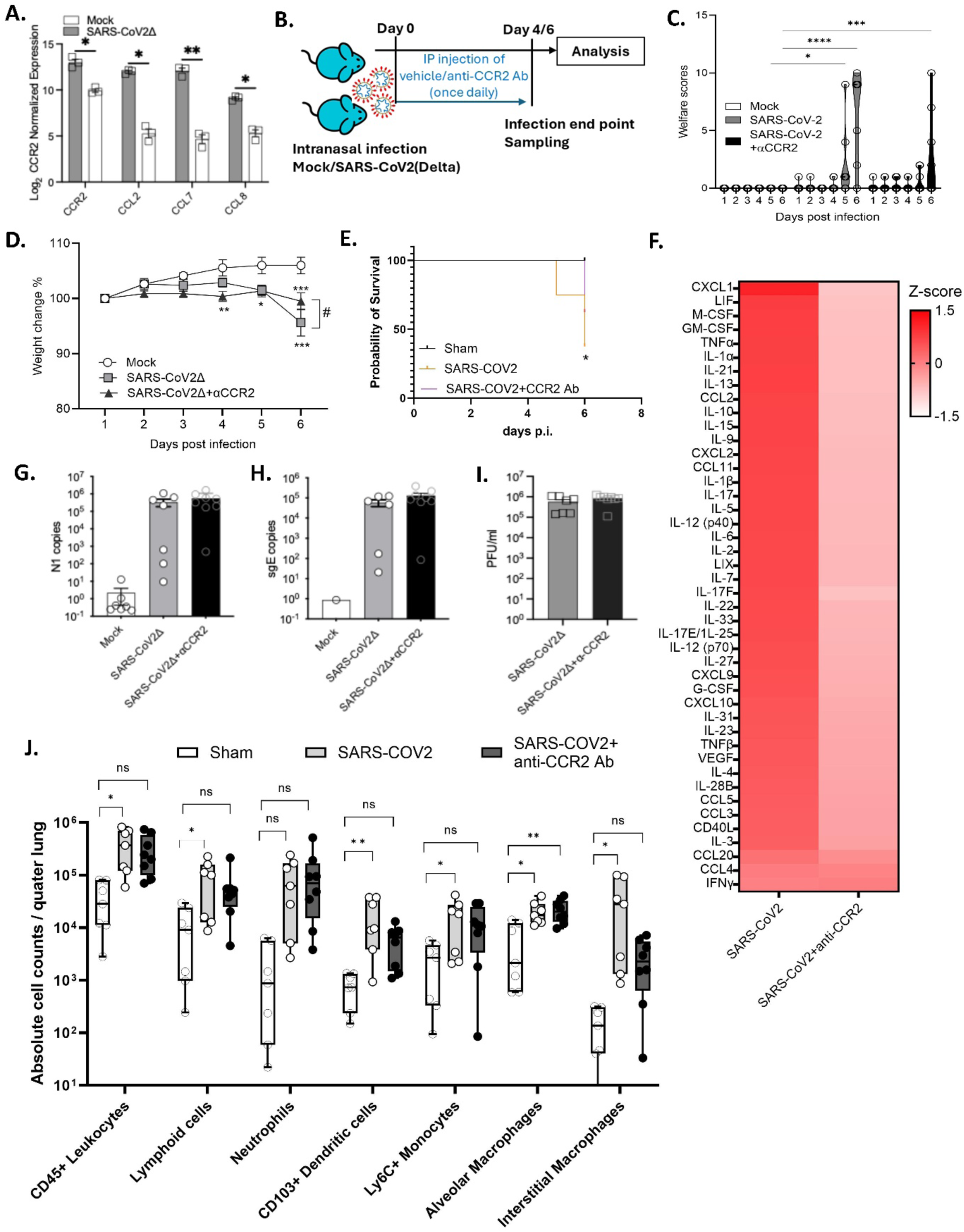
Anti-CCR2 treatment in SARS-CoV2Δ infected K18-hACE2 mice ablates the hyper-inflammatory response. **A**. normalized gene expression of CCR2 and its ligands from transcriptomic analysis of control and SARS-CoV2Δ infected lung. Data represented as mean±SEM. (* p≤0.05, ** p≤0.01; unpaired t-test). **B**. Experimental design of SARS-CoV2Δ experimental infections and daily treatments with MC-21 anti-CCR2 antibody. **C.** Total welfare scores and **D**. percentage weight change in mice following 6dpi of SARS-CoV2Δ infection and anti-CCR2 treatment. Data represented as mean±SEM (* p≤0.05, ** p≤0.01, *** p≤0.001, **** p≤0.0001 vs mock; p≤0.05 vs SARS-CoV2Δ; Two-way ANOVA and Dunnett’s post-hoc test, n=7-8/ group). **E.** Cumulative survival probability of 23 mice over 6 days of infection (p≤0.05 = *; n=7-8 / group) **F**. Heatmap showing z-scores of cytokines and chemokines in lung lysate quantified by a multiplex immunoassay. **G**. N1 copies and **H.** sgE viral RNA copies from lung quantified using one-step qPCR. Data represented as mean±SEM. **I.** Viral yields quantification from lung tissues by plaque assay. Data represented as mean±SEM. **J.** Box plots showing absolute cell counts of different immune cell sub-types in the mock, SARS-CoV2Δ infected and anti-CCR2 treated mice lungs. Each dot represents an individual animal. (p≤0.05 = *; p≤0.01 = **; one way-ANOVA with Dunnett’s post-hoc test).

To decipher the role of CCR2-dependent cell recruitment to the lung during inflammatory response to infection, and relevance to pathogenesis of severe disease, a group of infected mice were treated with anti-CCR2 antibody (MC-21) once daily until end of the experiment. (Fig. 3B). In our prior studies, this antibody treatment regimen has been effective in mediating profound depletion of CCR2+ inflammatory monocytes from circulation and lymphatic tissues in B.6 mice^20^. Both groups of SARS-CoV2Δ infected mice showed a significant weight loss from day 4 onwards, but the anti-CCR2 treatment groups showed significantly decreased weight loss at day 6 and better welfare scores at day 5&6 as compared to infection alone (Fig. 3C-D). Cumulative data of 23 mice assessed for 6 days post-infection showed significant increase in the survival probability of animals treated with anti-CCR2 antibody, (defined by attaining a humane endpoint theshold of clinical severity and/or weight loss of ≥10%, Fig.3E; 37.5% survival probability in vehicle treated vs 62.5% survival probability in anti-CCR2 treated, p<0.05, log-rank analysis for survival). Despite improved clinical scores, there were no differences in lung viral loads or yields between the infected and anti-CCR2 antibody treated groups following 6dpi (Fig.3 G-I).

Further, to evaluate the impact of anti-CCR2 treatment on pneumonitis in the lung during SARS-COV2Δ infection, we carried out immunophenotyping analysis of pulmonary cells 6d post infection in presence or absence of anti-CCR2 antibody treatment. As with day 5 immunophenotyping (Fig 1), we recorded a significant increase in the expansion of lymphoid cells (MHC-II^+^ CD11b^low^), inflammatory monocytes (CD11b^+^, Ly6C^+^), alveolar macrophages (SiglecF^high^, CD11c^+^, CD64^+^), dendritic cells (CD11b^−^, CD103^+^, CD24^+^, CD11c^+^), and interstitial macrophages (CD11b^+^, CD64^+^) into the infected lung (Fig 3J, p<0.05 vs mock, one way-ANOVA with Dunnett’s post hoc multiple-comparison test). Anti-CCR2 antibody treatment decreased the recruitment of inflammatory monocytes (CD11b^+^, Ly6C^+^). In addition to this, anti-CCR2 treatment also decreased the overall abundance of leukocytes (CD45+), lymphoid cells (MHC-II^+^ CD11b^low^), dendritic cells (CD11b^−^ CD103^+^ CD24^+^ CD11c^+^) and interstitial macrophages (CD11b^+^, CD64^+^) in infected lungs (Fig. 3J, p>0.05 vs mock, one way-ANOVA with Dunnett’s post hoc multiple-comparison test). However, the treatment had no effect on alveolar macrophage (SiglecF ^high^ CD11c^+^ CD64^+^) expansion within infected lungs (Fig. 3J). Concomitantly, the depletion of CCR2 cells using MC-21 antibody treatment led to a significant ablation of almost all 44 cytokine and chemokines quantified by multiplex immunoassay (Fig.2F). In particular, we identified a significant decrease in the levels of pro-inflammatory cytokines: IL-1α, IL-1β, IL-2, IL-5, IL-6, IL-7, TNF-α, IL-17F, GM-CSF and chemokines: CCL2, CXCL2, CCL11, CXCL1, and CXCL6 following the treatment (Fig.2F and SFig.6, p<0.05, unpaired t-test) (Fig.3J and SFig.6). Taken together these data suggest an expansive role of the CCR2-specific inflammatory response in SARS-CoV2 pulmonary hyperinflammation and severe COVID-19.

## Discussion

Following the global emergence of the COVID-19 global pandemic, there has been accumulating evidence supporting the use of various rodent preclinical systems including K18hACE-2 transgenic mice, as appropriate models of severe CoV2 pneumonitis^11–17^. SARS-CoV2 experimental infection in K18-hACE2 mice results in oedema-associated acute lung injury similar to that seen in COVID-19 patients, including coagulopathic and histological aspects of CoV-2 induced ARDS^40^. In this study, we confirm that the K18-hACE2 mouse model suitably recapitulates severe pulmonary disease after low-dose intranasal infection with the aggressive SARS-CoV2 delta VoC. Concomitant with viral gene replication in the lung, we observed welfare deterioration and weight loss from day 3 onwards and a decreased survival probability for the infected male mice from day 4. We recorded a cellular pneumonitis with increases in cell populations implicated in human severe lung disease, namely monocytes, macrophage populations and neutrophils.

Clinical data suggests gender-associated severity bias in COVID-19 patients. Although both males and females are equally susceptible to SARS-CoV2 infection, males experience higher severity and fatality^41–43^. Similar sex-based difference in lung viral load as well as disease severity has been observed following SARS-CoV2 infection in hACE2 mice^44–45^. In agreement with these reports, we also observed that male hACE2 mice were significantly more vulnerable to weight loss and reduced welfare following SARS-CoV2Δ inoculation. Male mice also showed relatively higher viral load in lungs compared to female mice. Whilst the basis behind this bias is not entirely clear, sex hormones influencing hyperinflammation, variability in ACE1/2 expression levels and dysregulated pro-coagulation responses has been reported in male versus female K18-hACE2 mice^44–45^.

We therefore focused our characterisations of cellular and molecular inflammatory disease processes occurring within more vulnerable male mice following SARS-CoV2Δ. Several clinical studies have undertaken in-depth transcriptomic analysis of severe COVID and ARDS, utilising lung biopsies collected from deceased patients, or cells from lung lavage following severe infection, to identify correlative markers of potential mechanistic significance in driving disease pathophysiology^46^. In this study, our pulmonary specific transcriptomic data from hACE2 mice recapitulated clinical biomarkers of COVID severe disease, including the IL-6 signalling pathway, TNF-α pathway, multiple chemokine pathways, and the type-1 interferon signalling pathway. Utilising cytokine / chemokine protein arrays, we corroborated transcriptomic analyses, whereby elevated pro-inflammatory cytokine and chemokine profiles were evident in SARS-CoV2Δ infected K18hACE2 mouse lungs, namely: IL-6, IFN-γ, CXCL-10, CCL2 and TNF-α, this correlates with clinical findings in COVID-19 patients^31–32,37–38^.

In a further validation of the hACE2 model to identify key signatures of inflammation leading to severe lung disease during SARS-CoV2Δ infection, we utilised the active metabolite of remdesivir, GS441524, to successfully modulate lung viral titres. We selected GS441524, the nucleoside active metabolite of remdesivir, as it has previously been demonstrated to successfully inhibit SARS-CoV2 in mice^17, 26–27^. Furthermore, remdesivir was the first antiviral to be granted emergency use authorization for treatment in human cases^25^. A three-day course of remdesivir in non-hospitalized COVID patients resulted in 87% lower risk of hospitalization or death (DOI: 10.1056/NEJMoa2116846). Although latter large-scale evaluations have cast doubt on the effectiveness of remdesivir, combining remdesivir with dexamethasone has shown to improve survival compared with dexamethasone alone^47–49^. Our data confirmed a median reduction in lung viral titres at 4dpi in GS441524-treated hACE2 mice. Concomitantly, treatment modified 44% of host response gene transcripts upregulated following infection, measured by NanoString. Further investigation of inflammatory protein levels in infected lungs identified several key signature chemokines and cytokines were significantly modified by GS441524, including CXCL-10, CCL2 and IL-6. In severe COVID, remdesivir treatment decreases the time to improve clinical symptoms after initiating treatment^50^. A recent study reported that remdesivir also significantly reduced circulating levels of IL-6, IL-10, IFN-α and CXCL10 in COVID patients after 5 days treatment^51^. Whether the anti-inflammatory impact of remdesivir during COVID is solely via modification of viral replication, or potentially via an additional anti-inflammatory mode of action of the drug, remains to be resolved. For example, remdesivir can reduce inflammation in ulcerative colitis-propelled gut inflammation in rodent models by enhancing the SIRT6/FoxC1 anti-inflammatory cascade, suggesting direct anti-inflammatory efficacy^52^.

Interestingly, we did not record an effect of GS441524 treatment on any of the major ISG genes that are upregulated in the lungs following SARS-COV2 infection. A previous study has shown that increased expression of 4 ISG genes, namely, *MX1, MX2, ISG15*, and *OAS1,* were associated with a higher viral load^53^. Potential uncoupling of inflammatory versus anti-viral immune pathways following remdesivir treatment, whereby the latter process remains intact, may contribute to overall efficacy and protection of severe disease by allowing host-anti-viral responses to reduce viral load. Additional longitudinal studies of ISGs expression following GS441524 treatment, and loss of function studies of ISG genes may resolve whether ISG pathways contribute to the remdesivir nucleoside analogue mode of action.

Treatment with dexamethasone, a potent glucocorticoid, that can blanket-suppress pro-inflammatory signaling pathways via transcription suppression, significantly lowers incidence of death in hospitalized COVID patients receiving invasive or non-invasive respiratory support and has become the mainstay treatment of severe ARDS due to SARS-CoV2^7^. However, deleterious sequalae of off-target effects post long-term 6mg/day dosages of dexamethasone include skin thinning, weight gain, osteoporosis, hypertension and diabetes^54^. More selectively targeting treatments of critical inflammatory pathways would be desirable to overcome side effects of glucocorticoid therapy. As such, IL-6 pathway inhibitors, (i.e. tocilizumab and sarilumab), and auto-immune pathway inhibitors (i.e. baricitinib) have been clinically investigated and proven to result in reduced risk of death and higher probability of improvement in hospitalized patients with severe COVID-19^8–9,55^. Among significantly upregulated genes found in SARS-CoV2Δ infected lungs in this study, the CCL2-CCR2 signaling pathway was predominant. Following SARS-CoV2Δ inoculation, CCR2 and its ligands: CCL2, CCL7 and CCL8, were some of the most-fold induced gene transcripts in infected lungs. Protein immunoassay corroborated CCL2 as one of the most highly upregulated molecules in the lung following SARS-CoV2Δ infection. Prior studies have identified CCR2-CCL2 signaling as crucial for the initial infiltration of inflammatory monocytes to the lung and for the expansion of monocyte-derived cells which act to limit SARS-CoV2 infection^38–39^. However, uncontrolled CCL2 expression, under the excessive influence of CCL2 inducers like IL-6, TNF-α, IFN-γ, and increased CCR2 signaling has been implicated as a central mediator of cytokine storm, hyper-inflammation and tissue damage causing severe disease. Additionally, the 3p21.31 locus, which controls the chemotactic receptor expression in monocytes and macrophages, is strongly associated with increased risk of morbidity and mortality^56^. Furthermore, a positive association between MCP-1-A2518 G (CCL2) gene variants with the severity of COVID-19 has been found^57^. Here we demonstrate significant therapeutic benefit of targeting the CCL2-CCR2 axis by application of the anti-CCR2 antibody MC-21^18–20^ which profoundly and temporarily depletes CCR2-bearing immune cells, particularly inflammatory monocytes, in blood and various tissue sites of inflammation. We demonstrate that following daily administration of MC-21 for six days during SARS-CoV2Δ infection, we successfully modulated pulmonary monocytes and, concomitantly, interstitial macrophages (potentially via reducing the pulmonary pool of monocytes for macrophage differentiation). MC-21 antibody treatment has been previously shown to similarly inhibit the influx of inflammatory monocytes, interstitial macrophages and differentiated T-cells to the lung in a mouse model of pneumococcal pneumonia, whereas it had no effect on alveolar macrophages^18^. Interstitial macrophages are found in the lung parenchyma and are also implicated in SARS-CoV2 infection as a prominent site of viral. Interstitial macrophages, in parallel with CCR2+ infiltrating monocytes, are major source of inflammatory cytokines and play key role in orchestrating airway inflammation following SARS-CoV2 infection^58–59^.

Anti-CCR2 treatment profoundly reduced cytokine-storm in the lung, which implicates recruitment of CCR2-bearing cells as central to this disease phenomenon during SARS-CoV2 infection. The dual impact of reduced cellular pneumonitis and reduced cytokine levels in the lung following MC-21 anti-CCR2 treatments was an improvement in welfare and protection from acute weight loss. A prior preclinical study, utilising the SARS-CoV2 Beta (β) VoC strain B.1.351 in K18hACE2 mice, has also demonstrated the benefit of manipulating the CCL2/CCR2 axis to treat SARS-CoV2 infections by use of a CCL2 antibody. CCL2 antibody treatment in K-18 hACE2 transgenic mice following the experimental infection of SARS-CoV2Δ resulted in improved welfare and delayed mortality^60^. Further, a dual chemokine receptor CCR2/CCR5 inhibitor, cenicriviroc, has been found to inhibit SARS-CoV2 replication *in vitro* and reduced virus-induced cell destruction^61^, however, separate randomised double-blind, phase II trials that evaluated cenicriviroc efficacy in COVID patients showed no difference in the recovery time of hospitalized patients^62–63^. While targeting of CCL2/CCR2 signalling has been used to modulate hyper-inflammation in other inflammatory diseases,^64–65^ further detailed clinical studies are necessary to critically evaluate the efficacy of targeting the CCR2/CCL2 axis in COVID-like viral ARDS syndromes.

In conclusion, we demonstrate, in a humanised mouse model of SARS-CoV2*Δ,* a central role for CCR2 and its ligands during hyper-immune mediated severe lung disease, which is modifiable both by remdesivir nucleoside analogue treatment or by experimental targeting of CCR2 cells. With an expanding number of selective pharmacological and biological therapies becoming available, our data supports the need for further detailed pre-clinical and clinical evaluations to identify new safe and effective inhibitors or small molecules targeting the CCR2/CCL2 signalling network as a promising novel approach to treat or prevent respiratory distress in COVID patients, in tandem with direct acting anti-viral therapeutics.

## Acknowledgements

This work was funded by an MRC Confidence in Concept grant to LSTM (MC_PC_19045).

## Notes

### Competing Interest Statement

The authors have declared no competing interest.

