## Supplementary file for "The CCR2 inflammatory pathway is a target for improving severe disease and pulmonary inflammation in experimental COVID-19"

**Supplementary data:**

| Stage | Marker | Fluorophore | Manufacturer | Clone | Dilution |
|---|---|---|---|---|---|
| Viability Dye | Live/Dead | efluor506 | Invitrogen | | 1:500 |
| Surface Antibody Cocktail | CCR2 | BV510 | BioLegend | SA203G11 | 1:150 |
| | Siglec-H | BV605 | BD | 440c | 1:150 |
| | Ly6G | BV650 | BioLegend | RB6-8C5 | 1:150 |
| | MerTK | BV711 | BD | 108928 | 1:150 |
| | Ly6C | BV785 | BioLegend | HK1.4 | 1:150 |
| | CD45 | FITC | BioLegend | 30-F11 | 1:150 |
| | CD11c | PE | eBiosciences | N418 | 1:150 |
| | CD11b | PerCP-Cy5.5 | eBiosciences | M1/70 | 1:150 |
| | CD24 | PECy5 | BioLegend | M1/69 | 1:150 |
| | CD103 | PE/Dazzle594 | BioLegend | QA17A24 | 1:150 |
| | F4/80 | APC-Fire750 | BioLegend | BM8 | 1:150 |
| | CD64 | PE-Cy7 | BioLegend | X54-5/7.1 | 1:150 |
| | MHC-II | APC | eBiosciences | M5/114.15.2 | 1:150 |
| | CD206 | AlexaFluor647 | ABD Serotec | MR5D3 | 1:150 |
| | Siglec-F | AlexaFluor700 | Invitrogen | IRNM44N | 1:150 |
| T Cell markers | B220 | BV570 | BioLegend | RA3-6B2 | 1:150 |
| | CD3 | APC/Cy7 | BioLegend | 17A2 | 1:150 |
| | CD4 | BV421 | BioLegend | GK1.5 | 1:150 |
| | CD8a | SparkBlue550/AF532 | BioLegend | 53-6.7 | 1:150 |

*Table S1:Markers used for Flow Cytometry Analysis*

| Name | 5’ to 3’ Sequence |
|---|---|
| nCOV_N1 | |
| Probe | FAM/ACC CCG CAT TAC GTT TGG TGG ACC ZEN/IABkFQ |
| Primers | FW GAC CCC AAA ATC AGC GAA AT<br>RV TCT GGT TAC TGC CAG TTG AAT CTG |
| E_Sarbeco | |
| Probe | FAM/ACA CTA GCC ATC CTT ACT GCG CTT CG ZEN/IABkFQ |
| Primers | FW ACA GGT ACG TTA ATA GTT AAT AGC GT<br>RV ATA TTG CAG TAC GCA CAC A |
| 18s(mouse) | |
| Probe | HEX/TCA AAG ATT AAG CCA TGC ATG TCT AAG TAC GCA C ZEN /IABkFQ |
| Primers | FW AGC CAT TCG CAG TTT TGT AC<br>RV ACC TGG TTG ATC CTG CCA GGT AGC |

*Table S2: PCR primers used in the study.*

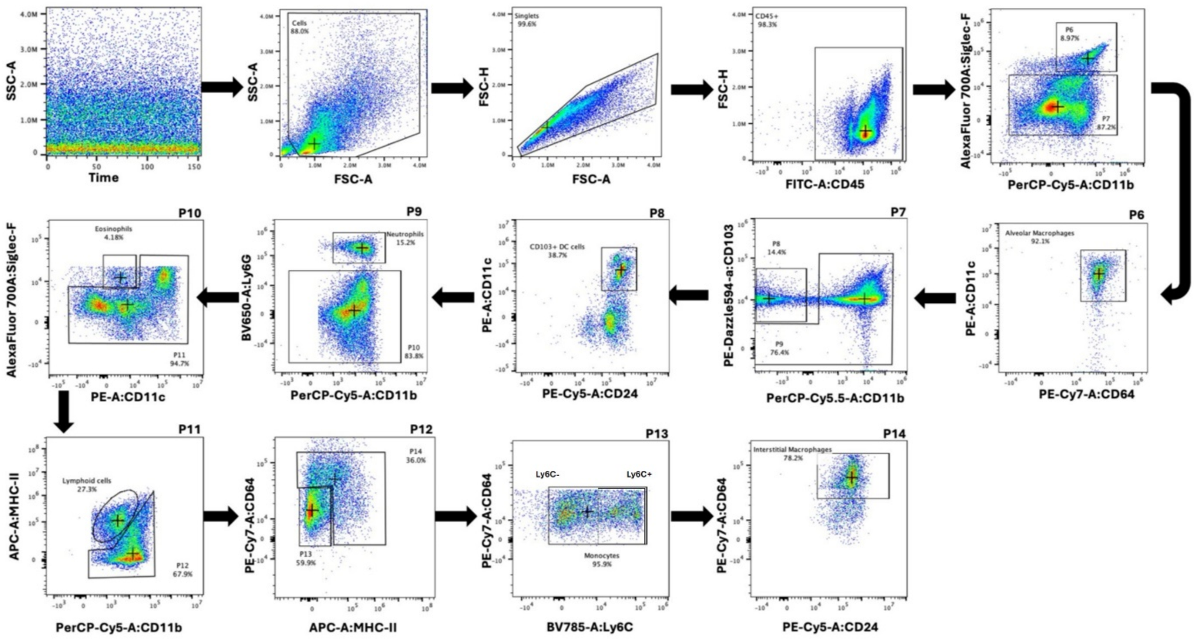


*Supplementary Figure 1: Gating strategy used in Flow cytometry (Misharin et al., 2013)*

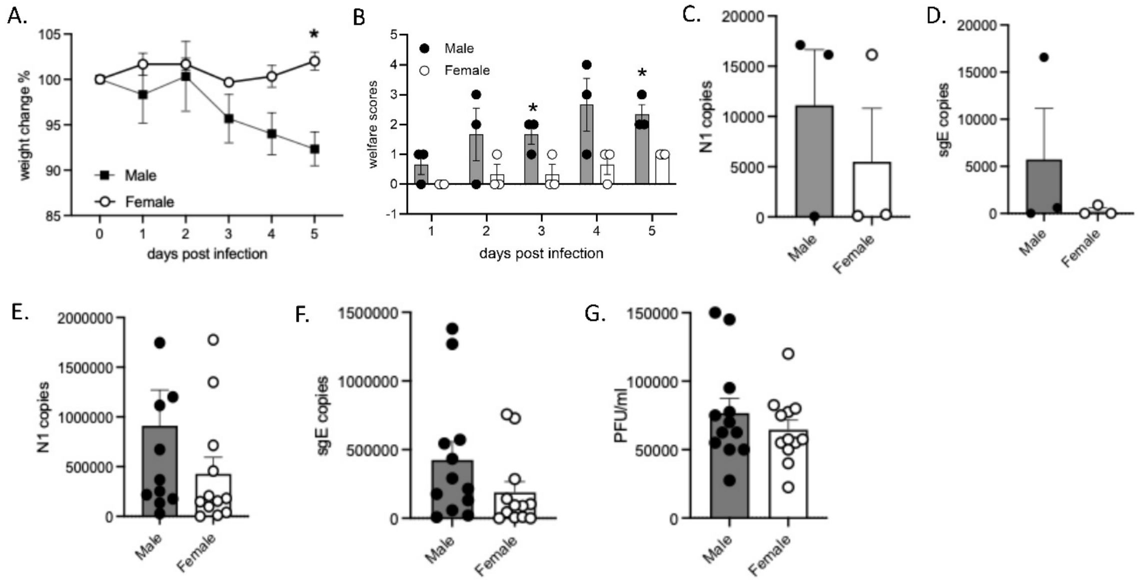


*Supplementary Figure 2: Male K18-hACE2 mice display more pronounced SARS-CoV-2 infection and clinical disease. (A) Weight changes in male and female mice following SARS-CoV2Δ infection. Data represented as Mean±SE (p≤0.05, multiple unpaired t-test) (B) Welfare scores in male and female mice following SARS-CoV2Δ infection. Data represented as Mean±SE (p≤0.05, multiple unpaired t-test). (C) One step qPCR quantification of N1 and (D) sgE RNA copies in lung tissue at 5 days post infection. Similar representation of differences in trends in viral quantification between male and female mice infected with the Wuhan variant over 4 days of infection. (E) N1, (F) sGE RNA copies and (G) Viral yields from lung tissues by plaque assay at 5 days post infection. Data represented as Mean±SD.*

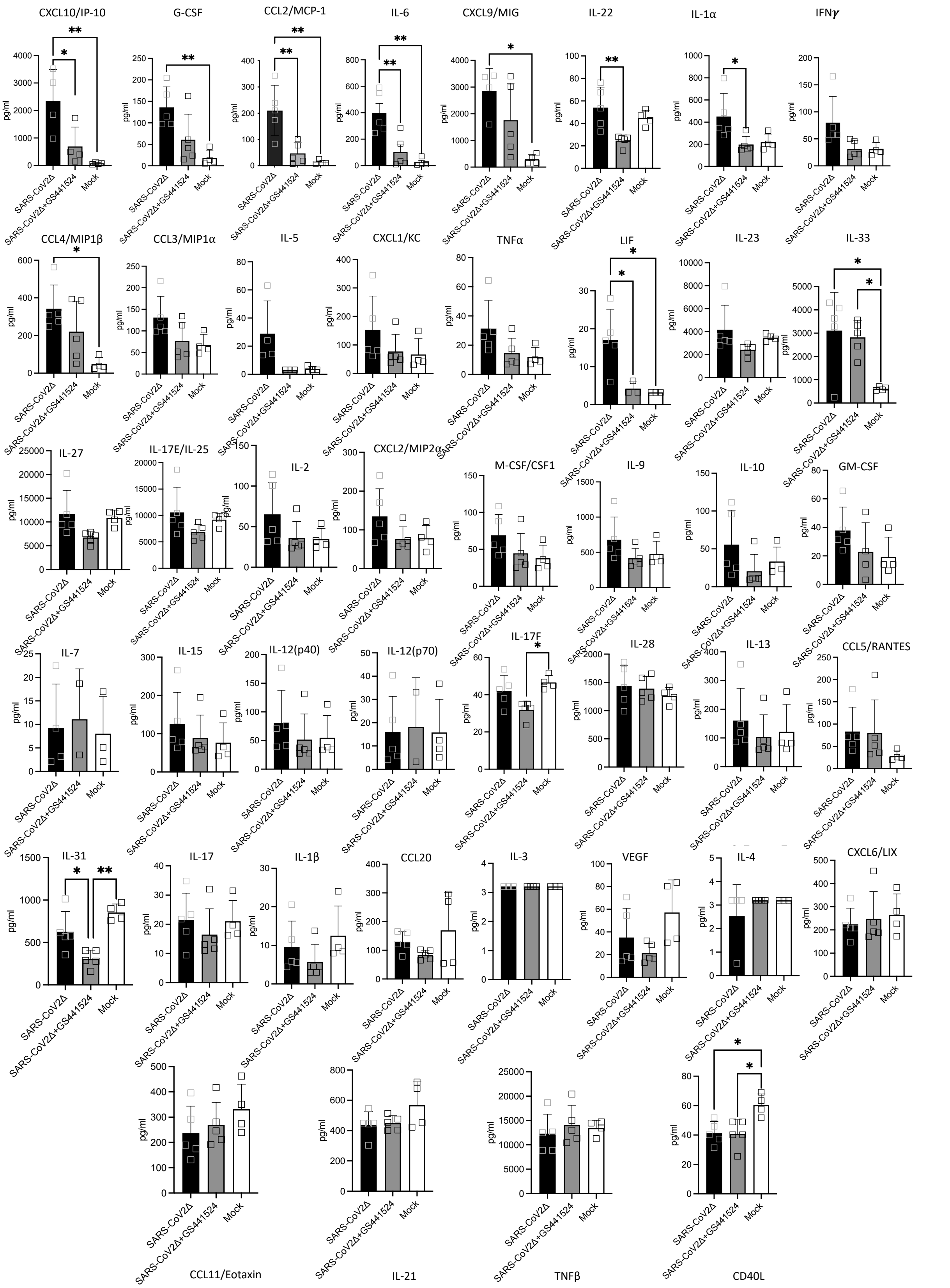


*Supplementary Figure 3:* ***Cytokine and chemokine expression in the lung tissues of SARS-CoV2infected mice as compared to those infected and treated with GS441524 over 4 days of infection*** *(as depicted in heatmap Fig 2E). Mice were infected intranasally with 2000PFU/ml of the Delta variant of SARS-CoV2 and monitored daily over 4 days. Lung tissue samples were then used to measure cytokines and chemokines using a 44-panel multiplex immunoassay. Data is presented as mean±SD and comparisons were made using a one-way ANOVA, * p<0.05, ** p<0.01 indicates significant difference (one-way ANOVA,Tukey's post-hoc test) .*

A.

**Gene Set Analysis**

| Term | Significance Score |
|---|---|
| Cytosolic sensors of pathogen-associated DNA | 10.1072 |
| DDX58/IFIH1-mediated induction of interferon-alpha/beta | 9.4737 |
| Signaling by GPCR | 6.7303 |
| Immunoregulatory interactions between a Lymphoid and a non-Lymphoid cell | 6.7099 |
| Toll-like Receptor Cascades | 6.172 |
| TNFR2 non-canonical NF-kB pathway | 5.889 |
| Class I MHC mediated antigen processing & presentation | 5.5546 |
| Metabolism of RNA | 5.4735 |
| DNA Repair | 5.3065 |
| Interleukin-4 and Interleukin-13 signaling | 5.2096 |

B.

**BioPlanet**

| Term | p-Adj |
|---|---|
| Interferon signaling | 1.0e-06 |
| Interferon alpha/beta signaling | 5.0e-06 |
| Thymic stromal lymphopoietin (TSLP) pathway | 0.00375 |
| Immune system | 0.00718 |
| Immune system signaling by interferons, interleukins, prolactin, and growth hormones | 0.01020 |

C. RIG-I/MDA5 mediated induction of IFN α/β pathways

### of Genes in Target **19** — **19↑ 0↓**

| Gene | -9 — +9 | Log2 FC | p-Value |
|---|---|---|---|
| Isg15 | | 6.80641 | 1.97e-08 |
| Irf7 | | 6.05425 | 7.11e-08 |
| Ifih1 | | 4.15451 | 6.49e-06 |
| Dhx58 | | 4.06311 | 9.26e-07 |
| Ddx58 | | 2.99672 | 5.73e-07 |
| Ube2l6 | | 2.89625 | 3.89e-05 |
| Nlrc5 | | 2.12309 | 0.001957 |
| Casp8 | | 1.96982 | 4.99e-05 |
| Irf1 | | 1.87003 | 7.39e-06 |
| Rnf135 | | 1.69001 | 0.000741 |
| Nfkb2 | | 1.57863 | 8.44e-05 |
| Trim25 | | 1.50678 | 2.30e-05 |
| Nfkbia | | 1.09963 | 0.004524 |
| Traf6 | | 1.08033 | 0.002389 |
| Tank | | 1.04056 | 2.56e-05 |
| Ager | | 0.870218 | 0.020388 |
| Crebbp | | 0.855679 | 0.009292 |
| Atg12 | | 0.725393 | 0.002831 |
| Nfkb1 | | 0.591756 | 0.000383 |

D. Interferon α/β signaling

### of Genes in Target **28** — **28↑ 0↓**

| Gene | -9 — +9 | Log2 FC | p-Value |
|---|---|---|---|
| Oas3 | | 7.7968 | 1.92e-07 |
| Isg15 | | 6.80641 | 1.97e-08 |
| Irf7 | | 6.05425 | 7.11e-08 |
| Ifit3 | | 5.65592 | 1.16e-07 |
| Oasl1 | | 5.3862 | 3.78e-06 |
| Oas1a | | 4.95255 | 3.39e-09 |
| Ifit2 | | 4.70601 | 1.39e-07 |
| Gbp2 | | 4.20795 | 6.65e-06 |
| Oas2 | | 3.63 | 5.03e-06 |
| Stat2 | | 3.50461 | 1.10e-05 |
| Xaf1 | | 3.46743 | 9.93e-07 |
| Stat1 | | 2.92304 | 3.70e-07 |
| Psmb8 | | 2.83461 | 1.66e-06 |
| Ifitm3 | | 2.70926 | 2.28e-06 |
| Ifi35 | | 2.68656 | 6.38e-07 |
| Ptpn6 | | 2.67391 | 1.82e-05 |
| H2-K1 | | 2.50176 | 1.77e-06 |
| H2-T23 | | 2.49165 | 5.95e-05 |
| Adar | | 2.11091 | 6.47e-06 |
| Irf9 | | 2.05555 | 0.000902 |
| Socs3 | | 2.02374 | 0.000228 |
| H2-D1 | | 1.95817 | 7.86e-05 |
| Irf1 | | 1.87003 | 7.39e-06 |
| H2-Q2 | | 1.6935 | 4.39e-05 |
| Ifitm2 | | 1.29125 | 0.002829 |
| H2-M3 | | 1.09333 | 0.001809 |
| H2-Q10 | | 0.929297 | 0.026634 |
| Ifnar2 | | 0.702867 | 0.002908 |

E.

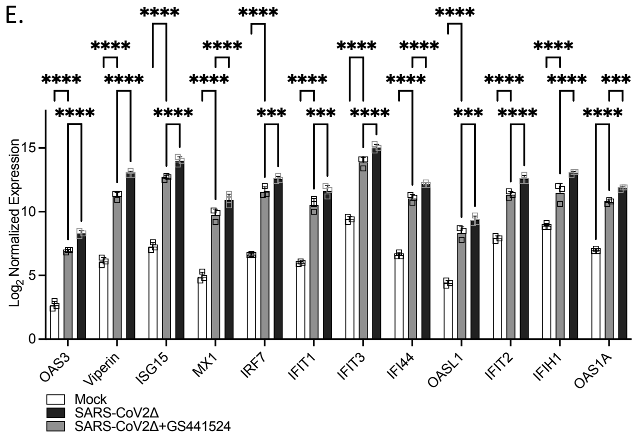


*Supplementary Figure 4* ***SARS-CoV-2 infection induces the expression of antiviral signaling genes in infected lung tissues and GS441524 treatment reduces the expression of some inflammatory markers*** *Mice were infected intranasally with 2000PFU/ml of the Delta variant of SARS-CoV2 and another group was additionally treated with GS441524 once daily and monitored daily over 4 days. Lung tissue samples were then used to measure gene expression by Nanostring analysis. A-B. Gene set analysis showing significant pathways when infected mice were compared to control mice. Expression of genes associated with C. Interferon $\alpha/\beta$ signaling and D. RIG-I/MDA5 mediated induction of IFN $\alpha/\beta$ pathways as determined by Nanostring analysis. E. Comparison of some interferon stimulated genes across mock, infected and infected and GS441524 treated mice. Data is presented as mean±SD.*

A.

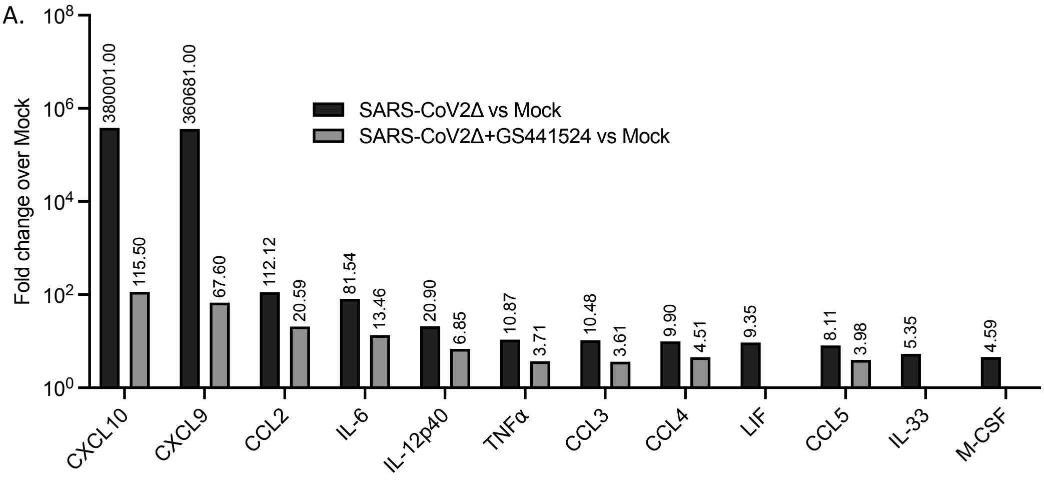


B. IL-6 signaling pathway

### of Genes in Target
6
6↑ 0↓

| Gene | Log2 FC | p-Value |
|---|---|---|
| Il6 | 6.34941 | 5.74e-06 |
| Socs3 | 2.02374 | 0.000228 |
| Stat3 | 1.33511 | 0.000191 |
| Il11ra1/2 | 1.18258 | 0.007461 |
| Jak2 | 0.997187 | 4.65e-05 |
| Il6ra | 0.755373 | 0.023489 |

C. Chemokine IL-8 signaling pathway

### of Genes in Target
16
16↑ 0↓

| Gene | Log2 FC | p-Value |
|---|---|---|
| Cxcl11 | 8.59495 | 2.99e-05 |
| Cxcl10 | 8.56986 | 2.09e-07 |
| Cxcl9 | 8.49458 | 3.87e-07 |
| Ccl7 | 7.74353 | 1.24e-06 |
| Ccl2 | 6.80889 | 9.49e-07 |
| Ccl12 | 4.56694 | 3.54e-06 |
| Xcl1 | 4.30212 | 1.29e-05 |
| Ccl8 | 3.90597 | 5.97e-06 |
| Cxcl13 | 3.43331 | 4.58e-05 |
| Ccl3 | 3.38904 | 1.62e-05 |
| Ccl4 | 3.30778 | 3.21e-05 |
| Ccl5 | 3.02002 | 2.09e-06 |
| Ccl19 | 2.9605 | 8.03e-06 |
| Ccl9 | 2.86598 | 1.16e-05 |
| Cxcl15 | 2.3783 | 0.000309 |
| Ccl11 | 2.10951 | 0.000185 |

D. Chemokine signaling pathway

### of Genes in Target
34
31↑ 3↓

| Gene | Log2 FC | p-Value |
|---|---|---|
| Cxcl10 | 8.56986 | 2.09e-07 |
| Ccl2 | 6.80889 | 9.49e-07 |
| Ccl12 | 4.56694 | 3.54e-06 |
| Xcl1 | 4.30212 | 1.29e-05 |
| Ccr5 | 3.72651 | 1.03e-06 |
| Stat2 | 3.50461 | 1.10e-05 |
| Cxcl13 | 3.43331 | 4.58e-05 |
| Ccl3 | 3.38904 | 1.62e-05 |
| Ccl4 | 3.30778 | 3.21e-05 |
| Ccr9 | 3.16599 | 5.21e-06 |
| Ccr2 | 3.10801 | 0.000167 |
| Ccl5 | 3.02002 | 2.09e-06 |
| Ccl19 | 2.9605 | 8.03e-06 |
| Stat1 | 2.92304 | 3.70e-07 |
| Hck | 2.51239 | 1.22e-05 |
| Ccl11 | 2.10951 | 0.000185 |
| Cxcl16 | 2.04872 | 0.000139 |
| Was | 1.87859 | 0.000177 |
| Ccr1 | 1.80222 | 0.000701 |
| Prkcd | 1.36501 | 2.36e-05 |
| Stat3 | 1.33511 | 0.000191 |
| Ccr7 | 1.09991 | 0.007166 |
| Nfkbia | 1.09963 | 0.004524 |
| Fgr | 1.01649 | 0.001756 |
| Jak2 | 0.997187 | 4.65e-05 |
| Ncf1 | 0.950596 | 0.017718 |
| Gsk3b | 0.910468 | 0.014087 |
| Pik3ca | -0.856804 | 9.18e-06 |
| Arrb2 | 0.792218 | 0.005602 |
| Raf1 | -0.639153 | 0.007708 |
| Rac2 | 0.63894 | 0.016304 |
| Mapk1 | 0.628917 | 0.007649 |
| Cxcr4 | -0.615762 | 0.016149 |
| Nfkb1 | 0.591756 | 0.000383 |

E. TNFα signaling pathway

### of Genes in Target
37
37↑ 0↓

| Gene | Log2 FC | p-Value |
|---|---|---|
| Cxcl10 | 8.56986 | 2.09e-07 |
| Il6 | 6.34941 | 5.74e-06 |
| Il1rn | 5.32657 | 3.92e-07 |
| Ccl12 | 4.56694 | 3.54e-06 |
| Il12b | 4.38543 | 4.16e-05 |
| Tnf | 3.44256 | 2.79e-05 |
| Ccl4 | 3.30778 | 3.21e-05 |
| Il2ra | 3.12985 | 1.49e-05 |
| Ccl5 | 3.02002 | 2.09e-06 |
| Cd69 | 2.62776 | 9.12e-05 |
| Vcam1 | 2.21452 | 1.42e-05 |
| Csf1 | 2.19838 | 2.55e-05 |
| Ripk2 | 2.16037 | 3.47e-05 |
| Ccl11 | 2.10951 | 0.000185 |
| Tnfsf10 | 1.94569 | 0.001712 |
| Cd44 | 1.89757 | 1.64e-05 |
| Lcp2 | 1.87611 | 1.82e-05 |
| Irf1 | 1.87003 | 7.39e-06 |
| Ccr1 | 1.80222 | 0.000701 |
| Csf2rb | 1.76706 | 2.38e-05 |
| Relb | 1.73266 | 4.93e-05 |
| Nfkb2 | 1.57863 | 8.44e-05 |
| Mapk8 | 1.54741 | 8.47e-05 |
| Il1r2 | 1.42183 | 0.029763 |
| Il7 | 1.2649 | 0.001186 |
| Il1b | 1.2209 | 0.011185 |
| Il1a | 1.20924 | 0.006230 |
| Cd40 | 1.20076 | 0.000348 |
| Ackr3 | 1.18086 | 0.003462 |
| Nfkbia | 1.09963 | 0.004524 |
| Il18r1 | 0.91271 | 0.015500 |
| Sod2 | 0.871896 | 0.000623 |
| Cmklr1 | 0.871662 | 0.008350 |
| Tgfb2 | 0.755859 | 0.012518 |
| Ifnar2 | 0.702867 | 0.002908 |
| Cflar | 0.673895 | 0.014068 |
| Ifngr2 | 0.640396 | 0.002452 |

F. Immune System

### of Genes in Target

172 166↑ 6↓

| Gene | Log2 FC | p-Value |
|---|---|---|
| Oas3 | 7.7968 | 1.92e-07 |
| Isg15 | 6.80641 | 1.97e-08 |
| Il6 | 6.34941 | 5.74e-06 |
| Irf7 | 6.05425 | 7.11e-08 |
| Ifit3 | 5.65592 | 1.16e-07 |
| Fcgr1 | 5.53359 | 9.90e-08 |
| Zbp1 | 5.4475 | 3.83e-08 |
| Oasl1 | 5.3862 | 3.78e-06 |
| Il1rn | 5.32657 | 3.92e-07 |
| Oas1a | 4.95255 | 3.39e-09 |
| Ifit2 | 4.70601 | 1.39e-07 |
| Gbp2 | 4.20795 | 6.65e-06 |
| Ifih1 | 4.15451 | 6.49e-06 |
| Tlr9 | 4.13559 | 3.67e-07 |
| Gbp5 | 4.11553 | 6.55e-06 |
| Dhx58 | 4.06311 | 9.26e-07 |
| Cd86 | 4.00459 | 3.31e-05 |
| C2 | 3.73704 | 2.37e-05 |
| Eif2ak2 | 3.72549 | 1.22e-06 |
| Gbp3 | 3.68333 | 3.87e-06 |
| Oas2 | 3.63 | 5.03e-06 |
| Cd274 | 3.5776 | 1.93e-06 |
| Stat2 | 3.50461 | 1.10e-05 |
| Xaf1 | 3.46743 | 9.93e-07 |
| Tlr7 | 3.46641 | 2.82e-05 |
| Psmb9 | 3.29069 | 4.24e-05 |
| Il2ra | 3.12985 | 1.49e-05 |
| Ctss | 3.11773 | 1.71e-06 |
| Ccr2 | 3.10801 | 0.000167 |
| Ddx58 | 2.99672 | 5.73e-07 |
| Lgals3 | 2.99238 | 1.15e-05 |
| Dtx3l | 2.93344 | 1.35e-05 |
| Stat1 | 2.92304 | 3.70e-07 |
| Ube2l6 | 2.89625 | 3.89e-05 |
| Psmb8 | 2.83461 | 1.66e-06 |
| Ifitm3 | 2.70926 | 2.28e-06 |
| Ifi35 | 2.68656 | 6.38e-07 |
| Ptpn6 | 2.67391 | 1.82e-05 |
| Pycard | 2.63985 | 2.09e-06 |
| Mefv | 2.5903 | 7.18e-05 |
| Psmb10 | 2.57913 | 1.66e-05 |
| Il2rb | 2.55471 | 8.04e-06 |
| Fcgr4 | 2.51858 | 3.16e-05 |
| Hck | 2.51239 | 1.22e-05 |
| H2-K1 | 2.50176 | 1.77e-06 |
| H2-T23 | 2.49165 | 5.95e-05 |
| Tap1 | 2.43051 | 6.18e-08 |
| Klrc1 | 2.34347 | 0.000147 |
| Il3ra | 2.31757 | 0.000102 |
| Vcam1 | 2.21452 | 1.42e-05 |
| Ripk2 | 2.16037 | 3.47e-05 |
| Nlrc5 | 2.12309 | 0.001957 |
| Adar | 2.11091 | 6.47e-06 |
| Hcst | 2.07433 | 0.000237 |
| Itgal | 2.07262 | 2.62e-05 |
| Cd80 | 2.0724 | 4.49e-06 |
| Klrd1 | 2.05628 | 1.19e-05 |
| Irf9 | 2.05555 | 0.000902 |
| Tlr2 | 2.04291 | 0.000297 |
| Tlr8 | 2.04014 | 0.000217 |
| Socs3 | 2.02374 | 0.000228 |
| Tap2 | 2.02116 | 0.002160 |
| Il2rg | 2.00401 | 0.000138 |
| Tlr6 | 2 | 9.94e-05 |
| Pirb | 1.99552 | 9.29e-05 |
| Lilra6 | 1.98457 | 0.000543 |
| Cd247 | 1.97525 | 4.73e-05 |
| Cd3e | 1.97421 | 0.000250 |
| Casp8 | 1.96982 | 4.99e-05 |
| H2-D1 | 1.95817 | 7.86e-05 |
| Cd44 | 1.89757 | 1.64e-05 |
| Was | 1.87859 | 0.000177 |
| Lcp2 | 1.87611 | 1.82e-05 |
| Irf1 | 1.87003 | 7.39e-06 |
| Sell | 1.82774 | 0.000135 |
| Casp4 | 1.7761 | 3.05e-05 |
| Fbxo6 | 1.77486 | 0.000427 |
| Lck | 1.77052 | 7.96e-05 |
| Cd96 | 1.76988 | 0.000612 |
| Csf2rb | 1.76706 | 2.38e-05 |
| Creb1 | 1.73386 | 0.001547 |
| Pstpip1 | 1.72757 | 0.001008 |
| Ptprc | 1.70892 | 0.000201 |
| Amica1 | 1.69832 | 0.000193 |
| Ncf4 | 1.69553 | 0.001434 |
| H2-Q2 | 1.6935 | 4.39e-05 |
| Rnf135 | 1.69001 | 0.000741 |
| Aim2 | 1.66763 | 0.001132 |
| Pik3ap1 | 1.64342 | 0.000568 |
| Panx1 | 1.62274 | 2.81e-05 |
| Trim21 | 1.59151 | 0.000114 |
| Nfkb2 | 1.57863 | 8.44e-05 |
| Prkcq | 1.57631 | 0.000689 |
| Card11 | 1.56181 | 0.005579 |
| Icos | 1.55336 | 0.001387 |
| Mapk8 | 1.54741 | 8.47e-05 |
| Pdcd1lg2 | 1.54689 | 0.007734 |
| Cd28 | 1.52696 | 0.006140 |
| Plcg1 | -1.52582 | 0.001895 |
| Il18 | 1.52223 | 0.000476 |
| Trim25 | 1.50678 | 2.30e-05 |
| H2-Eb1 | 1.49868 | 0.012110 |
| Hsp90b1 | 1.43168 | 0.002291 |
| Nlrc4 | 1.42649 | 5.26e-06 |
| Il1r2 | 1.42183 | 0.029763 |
| Tlr4 | 1.41638 | 0.000623 |
| Prkcd | 1.36501 | 2.36e-05 |
| Cd8b1 | 1.34685 | 0.002436 |
| C3 | 1.33593 | 7.38e-05 |
| Stat3 | 1.33511 | 0.000191 |
| Ifitm2 | 1.29125 | 0.002829 |
| Il7 | 1.2649 | 0.001186 |
| Rel | 1.26219 | 9.15e-05 |
| Nlrp3 | 1.23879 | 0.014549 |
| Il1b | 1.2209 | 0.011185 |
| Casp1 | 1.21343 | 0.000488 |
| Il1a | 1.20924 | 0.006230 |
| Cd40 | 1.20076 | 0.000348 |
| Cd3g | 1.1831 | 0.000949 |
| Cd3d | 1.17383 | 0.000396 |
| Mapkapk2 | 1.16944 | 0.001609 |
| Ifngr1 | 1.16693 | 0.011523 |
| Itgb2 | 1.16 | 0.000257 |
| Il1rap | 1.11816 | 0.002165 |
| Nfkbia | 1.09963 | 0.004524 |
| H2-M3 | 1.09333 | 0.001809 |
| Tlr1 | 1.0842 | 0.011983 |
| Bcl2 | 1.08196 | 0.007504 |
| Traf6 | 1.08033 | 0.002389 |
| Fcgr2b | 1.06587 | 0.001317 |
| H2-Ab1 | 1.06219 | 0.024722 |
| Kpnb1 | 1.05858 | 0.000483 |
| Tank | 1.04056 | 2.56e-05 |
| Myd88 | 1.02266 | 0.002142 |
| Jak2 | 0.997187 | 4.65e-05 |
| Map3k3 | 0.988839 | 0.004273 |
| Ripk3 | 0.968636 | 0.012205 |
| Zap70 | 0.961244 | 0.002860 |
| Ncf1 | 0.950596 | 0.017718 |
| Map2k4 | 0.934412 | 0.000564 |
| Rps6ka3 | 0.933151 | 0.000281 |
| H2-Q10 | 0.929297 | 0.026634 |
| Ncf2 | 0.917477 | 0.010239 |
| Gsk3b | 0.910468 | 0.014087 |
| Icosl | 0.887481 | 0.012215 |
| Rab7 | 0.875688 | 0.001258 |
| Hc | -0.873637 | 0.000833 |
| Ager | 0.870218 | 0.020388 |
| Pik3ca | -0.856804 | 9.18e-06 |
| Crebbp | 0.855679 | 0.009292 |
| [illegible] | [illegible] | [illegible] |

*Supplementary Figure 5:* ***SARS-CoV-2 infection is characterized by expression of inflammatory genes and activation of associated pathways.*** *Mice were infected intranasally with 2000PFU/ml of the Delta variant of SARS-CoV2 and another group was additionally treated with GS441524 once daily and monitored daily over 5 days. Lung tissue samples were then used to measure gene expression by Nanostring analysis. A. Differences in fold change gene expressions over mock control mice between infected mice as compared to infected and GS44152 treated mice. Significantly upregulated genes in various important pathways determined by Nanostring analysis when infected mice were compared with controls B. IL-6 signaling. C. Chemokine IL-8 signaling pathway D. Chemokine signaling pathway E. TNF$\alpha$ signaling pathway and F. Immune system pathway.*

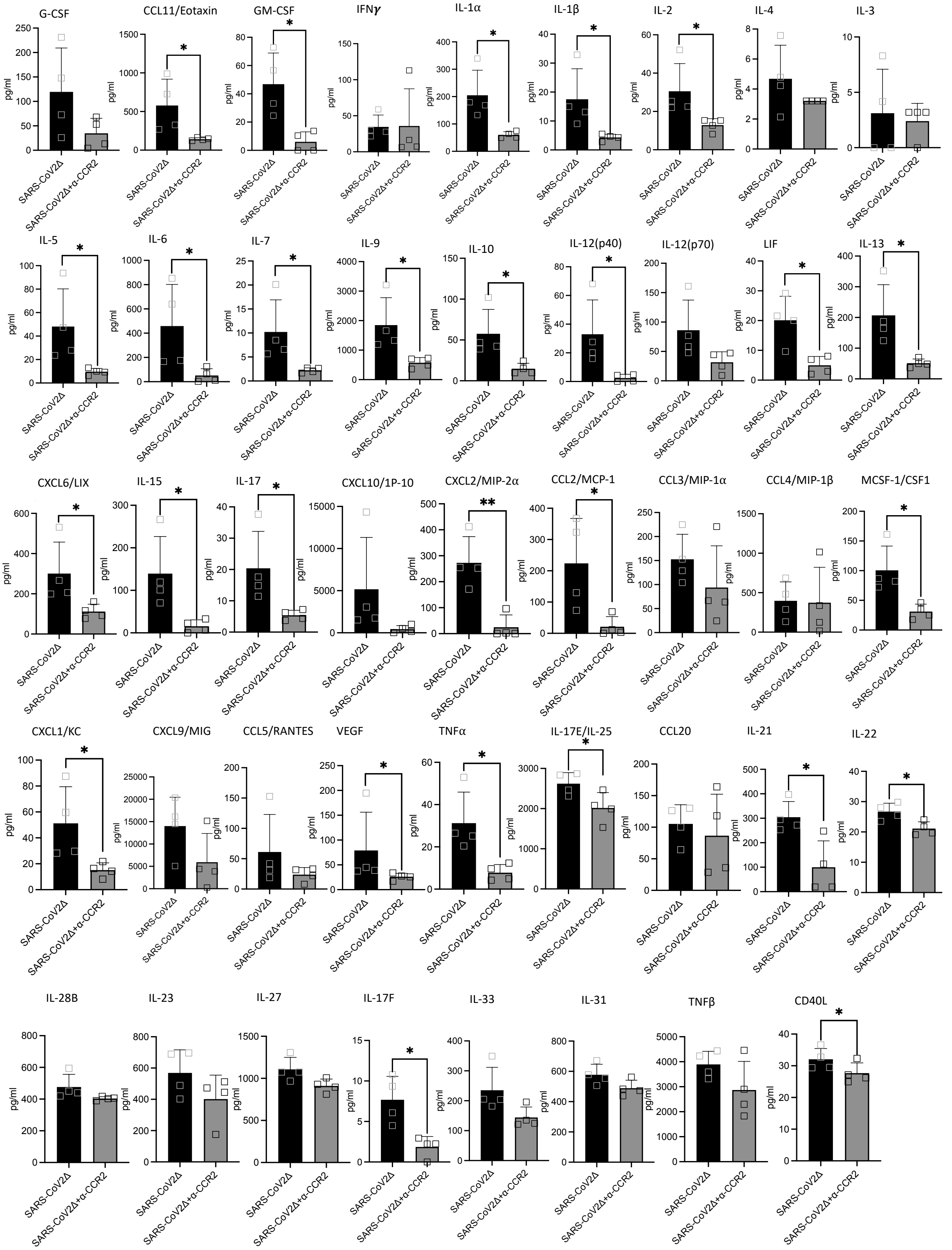


*Supplementary Figure 6:* ***Cytokine and chemokine expression in the lung tissues of SARS-CoV2 infected mice as compared to those infected and treated with anti-CCR2 antibody*** *(as depicted in heatmap Fig 3I). Mice were infected intranasally with 1000PFU/ml of the Delta variant of SARS-CoV2 and another group was additionally treated with anti-CCR2 antibody once daily and monitored daily over 5 days. Lung tissue samples were then used to measure cytokines and chemokines using a 44-panel multiplex immunoassay. Data is presented as mean±SD and comparisons were made using a unpaired t-test assuming unequal variance. * p<0.05, ** p<0.01 indicates significant difference.*
